# Ligand Binding Prediction using Protein Structure Graphs and Residual Graph Attention Networks

**DOI:** 10.1101/2022.04.27.489750

**Authors:** Mohit Pandey, Mariia Radaeva, Hazem Mslati, Olivia Garland, Michael Fernandez, Martin Ester, Artem Cherkasov

## Abstract

**Motivation:** Computational prediction of ligand-target interactions is a crucial part of modern drug discovery as it helps to bypass high costs and labor demands of in vitro and in vivo screening. As the wealth of bioactivity data accumulates, it provides opportunities for the development of deep learning (DL) models with increasing predictive powers. Conventionally, such models were either limited to the use of very simplified representations of proteins or ineffective voxelization of their 3D structures. Herein, we present the development of the PSG-BAR (Protein Structure Graph –Binding Affinity Regression) approach that utilizes 3D structural information of the proteins along with 2D graph representations of ligands. The method also introduces attention scores to selectively weight protein regions that are most important for ligand binding.

**Results:** The developed approach demonstrates the state-of-the-art performance on several binding affinity benchmarking datasets. The attention-based pooling of protein graphs enables identification of surface residues as critical residues for protein-ligand binding. Finally, we validate our model predictions against an experimental assay on a viral main protease (Mpro)– the hallmark target of SARS-CoV-2 coronavirus.

**Availability:** The code for PSG-BAR is made available at https://github.com/diamondspark/PSG-BAR

## 1 Introduction

Drug discovery field is constantly evolving to yield safe and potent drugs in the most time-, cost- and labor efficient way. Traditionally, wet-lab high-throughput screening (HTS) was performed with libraries of compounds tested against a target of interest (typically a protein) to identify potential drug candidates^1^. The drawbacks of such approach are in high cost and labor demand, further hindered by a modest (on average 0.03%) hit rate^2–4^. Thus, effective computational methods are urgently needed to help accelerating the drug discovery^2,5^.

For instance, virtual screening helps to narrow down the score of compounds for experimental validation from large chemically diverse databases thereby saving time and resources ^6,7^. The mostly used virtual screening tool, molecular docking relies mainly on physics-based or statistically derived scoring functions (SFs) to predict the binding affinity of ligands. Docking has achieved impressive advances in refining the drug discovery pipeline and helped identifying many potent and selective drug candidates^8,9^. Although docking is significantly faster than other virtual screening tools like quantum mechanics-based approaches and free-energy perturbation simulations, its speed is limited and does not allow covering the wealth of available chemical structures. For example, the new release of ZINC database^10^ containing over a billion molecules is impossible to screen with conventional tools and, thus, large chemical space remains unexplored. To overcome this issue, machine learning has been widely integrated into drug discovery pipelines. As such, DeepDocking^7^ integrates quantitative structure-activity relationship deep models with conventional docking software and achieves a 50-fold acceleration. Other machine learning (ML) models designed to predict drug-target interactions were shown to be capable of capturing the non-linear patterns and infer complex binding rules^11,12^. In fact, some of the traditional ML models and recently developed deep learning (DL) models outperformed classical virtual high throughput screening (vHTS) approaches in terms of speed and predictive performance^9^.

Several previous studies have leveraged DL-based models to predict drug-target interaction (DTI) and interaction binding affinity^13^. An efficient model should integrate both protein target and small molecule information and going beyond traditional pharmacophore descriptor^14^, 1D descriptors learned from protein sequences and ligand atoms respectively have gained significant traction^15–17^. Thus, DeepDTI approach predicted DTI using deep belief network model fed with extended-connectivity fingerprints for drug representations and sequence composition descriptor for proteins^18^. DeepDTA^19^ and DeepConv-DTI^20^ exploit a CNN architecture with protein and ligand sequences as inputs to predict binding affinity for Kinases.

However, complex patterns of protein-ligand interactions can only be captured with more realistic 3D structures and 3D and higher order descriptors that have been in the development over several decades by many groups around the world^21^, including our own^22–24^. Importantly, databases of experimentally derived protein crystal structures also grow by the day^25^, while powerful predictive tools such as AlphaFold^26^ further supplement the wealth of available protein structural data. To this end, recent studies more readily employ structural protein information alongside ligands. These methods demand faithful representation of biological targets – such as proteins – and drug compounds. Inspired by the advances in computer vision and 3D object recognition, researchers have explored 3DCNN for modeling protein structures^11,27,28^. As such, Jimenez et al (2018)^27^ employed 3D convolutional neural networks (3D-CNN) to predict ligand binding affinities. Jones et al (2021)^11^ developed a model to predict ligand-protein binding affinity using a fusion of 3D-CNN and spatial graph convolutional neural network.

While these approaches yield excellent predictive results, grids with atomic voids within the structure makes them susceptible to sparsity, memory bottleneck and inefficient computations. Furthermore, CNNs are sensitive to rotation and orientation. While this may be useful for different conformers of ligands, proteins are typically considered as fixed rigid bodies for binding. To overcome this expensive computation of 3DCNN for large protein structures alternative rotation invariant representations, namely graph neural networks, have been investigated.

In contrast to CNNs, graphs provide efficient and succinct representations of 3D structures. Several studies have reported superior performance on binding affinity prediction by representing proteins as a network of their amino acid residues^29–33^. For example, GraphBAR^31^ modelled protein ligand complexes as adjacency matrices at various spatial resolution of atoms of the binding site. Similarly, Lim et al (2019)^29^ propose a distance-aware representation of protein-ligand complex using two adjacency matrices. GEFA^30^ fuses 2 separate graphs (i) for protein using amino acid sequence and contact map and (ii) for the drug molecules from their constituent atoms. Their strategy to hard-mask the predicted unimportant regions is based on self-attention scores is restrictive as it relies strongly on the accuracy of their method’s binding region prediction.

Those methods benchmark their performance on PDBBind dataset where the active pocket structure of the protein and the protein-ligand complex is known. Obtaining such information is expensive and limited to relatively smaller datasets and hence the applicability of such methods is limited to a few carefully curated datasets. Furthermore, most of these methods fall short of reporting generalizability of their methods across targets and ligands.

To alleviate these limitations and challenges of classical docking as well as previous deep learning approaches, we introduce a graph-Siamese like network with attention-based graph pooling. Benefitting from the advantages of protein crystallographic structures, the developed method applies a graph attention network to the entire protein structures to predict several measures of ligand-protein binding affinities^34^. The graph-Siamese like network facilitates concurrent feature extraction and fusion of the protein and ligand graphs by the predictive model. The use of attention enables our model to locate the likely active residues of the protein for ligand binding, hence reducing the overhead of time-consuming manual preparation of protein structures necessary for docking. For the model evaluation, we employed publicly accessible databases - PDBbind^18^, BindingDB^19^, KIBA^33^, and DAVIS^35^, containing affinity measures for 9,777, 123,904, 23,732 and 41,142 complexes respectively (Table 1). We consequently retrieved protein structural information from PDB database and for KIBA dataset supplemented missing structures with those predicted by AlphaFold^26,36^. We demonstrate that the use of AlphaFold predicted structures enhances the performance of the model and that our apporach effectively predicts ligand-protein binding affinity and outperforms several existing baselines and circumvents expensive target site preparation needed for docking.

**Table 1.** Data statistics for datasets used for evaluation of PSG-BAR

| Dataset | Unique Targets | Unavailable PDBs | Unique Ligands | Unique Interactions |
| --- | --- | --- | --- | --- |
| PDBBind | 9,619 | 0 | 7,981 | 9,777 |
| KIBA | 467 | 181 | 2,356 | 124,374 |
| DAVIS | 442 | 139 | 68 | 20,604 |
| BindingDB | 1,038 | 100 | 13,222 | 41,142 |
| AID 1706 | 1 | 0 | 290,765 | 290,765 |

Our main contributions in this paper are

i. We propose an effective graph-Siamese like network, PSG-BAR, to simultaneously operate on structural graphs of proteins and molecular graphs of drugs for binding affinity prediction.
ii. We introduce an attention-based readout strategy to generate graph-level embedding for proteins. These attention scores are shown to correlate with some known physical properties of binding sites. We call this learning component *interaction attention module.*
iii. We investigate the benefit of adding Alphafold’s predicted protein structures to KIBA binding affinity dataset and demonstrate that it reduces variance and improves the performance metric of binding affinity prediction.

This paper is organized as follows: In section 2, we describe the datasets, their statistics, and the details of our proposed method. Section 3 features predictive results of our method on four benchmarking datasets and its application for several pertinent drug discovery tasks. In sections 4 and 5 we present a multi-view analysis of results and investigate the effect of architectural sub-components on overall performance of our method. Finally, section 6 discusses future directions and implications of this work.

## 2 Methods

### 2.1 Dataset

In this work we used four publicly available datasets on protein-ligand binding affinity including BindingDB^37^, PDBBind v2016^38^, KIBA^33^, and DAVIS^35^ that contain various binding affinity (Kd, Ki) datasets. First, we retrieved all protein-ligand pairs with associated dissociation constant (Kd) from BindingDB database^37^. Following data curation procedures such as removing duplicates, inorganic compounds, invalid SMILES, Kds outside of typical range and deriving associated structure files for each protein (PDBs with lowest resolution per protein target), the dataset was reduced to 41,142 protein-ligand pairs with 1,038 unique proteins and 13,222 unique ligands. The pKd values had a mean of 6 and standard deviation of 1.57. Similarly, we derived 13,478 unique interactions from PDBBind^38^. The 2016 dataset version of the PDBBind was used in this study in order to provide a consistent comparative analysis against a set of aforementioned studies. This PDBBind dataset includes three subsets: a core, a general, and a refined databases totaling 10,790 unique protein-ligand complexes which were narrowed down to 8,000 unique complexes after filtering on entries associated with Kd values. The Kd values were also transformed to pKd and had a mean STD of 6.35 and 1.89 respectively.

In addition to those values, we also retrieved data from the KIBA dataset^33^ which consists of a drug-target matrix with information on three major bioactivity types – IC50, Ki and Kd. Protein targets from the KIBA were labelled with UniProt IDs, which were then mapped to their respective PDB IDs to derive crystal structure. KIBA scores were transformed similarly to Shim et al. (2021)^39^ and He et al. (2017)^40^. In particular, drugs and targets with less than 10 interactions were filtered out, then KIBA scores were transformed to resemble pKd. Of the original 467 targets, after filtering and mapping to available PDB IDs, 286 targets remained. Of the 52,498 drugs, after filtered, there remained 2,356. This yielded 124,374 interaction pairs.

We also evaluated the proposed model on the DAVIS dataset^35^ containing 72 kinase inhibitors tested over 442 kinases which represent over 80% of the human catalytic protein kinome. The activity numbers distribution was similar to Bind-ingDB and PDBBind with a mean of 5.48 and std 0.928. Protein targets were labelled by name, so to retrieve corresponding PDB structures, the RCSB PDB^41^ Search API was used to perform a keyword search. Of the 442 proteins, 303 could be mapped to a PDB structure covering 68 of the initial 72 kinase inhibitors. In total, this yielded 20,604 drug-target interaction pairs.

To further evaluate applicability of our method for the SARS coronavirus (SARS-CoV-2) we attempted to predict inhibitors of the SARS-CoV-2 3C-like protease (3CLpro), also known as the main protease (Mpro). The bioassay record (AID 1706)^42^ by Scripps Research Institute provides PubChem Activity Score normalized to 100% observed primary inhibition. As suggested by authors of AID 1706, the activity was thresholded at 15 to get 444 inhibitors for 3CLpro along with 290,321 inactives. The protein structure corresponding to the crystal structure of SARS-CoV-2 Mpro in complex with an inhibitor N1 (1WOF^*43*^) was used.

For each of the datasets, certain targets and ligands were dropped on account of unavailability of the crystal structures and un-processability by RDKit respectively. Table 1 lists the counts of eventually used protein-ligand interaction pairs for each dataset.

### 2.2 Model Architecture

PSG-BAR receives two attributed graphs 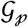 and 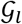 corresponding to the protein 3D structure and molecular graph of ligand’s 2D representation respectively (Figure 1). The construction of 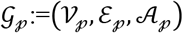 and 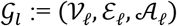 is as defined in section *Protein and Ligand graph construction method* (2.2). The model architecture follows an encoder-decoder paradigm with the encoder comprising of (i) protein encoder 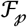 (ii) cross attention head, and (iii) ligand encoder 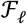. We describe the cross-attention mechanism in section *Interaction attention module* (section 2.6). The decoder is a multilayer feed forward neural network with LeakyReLU activation.

**Figure 1.**
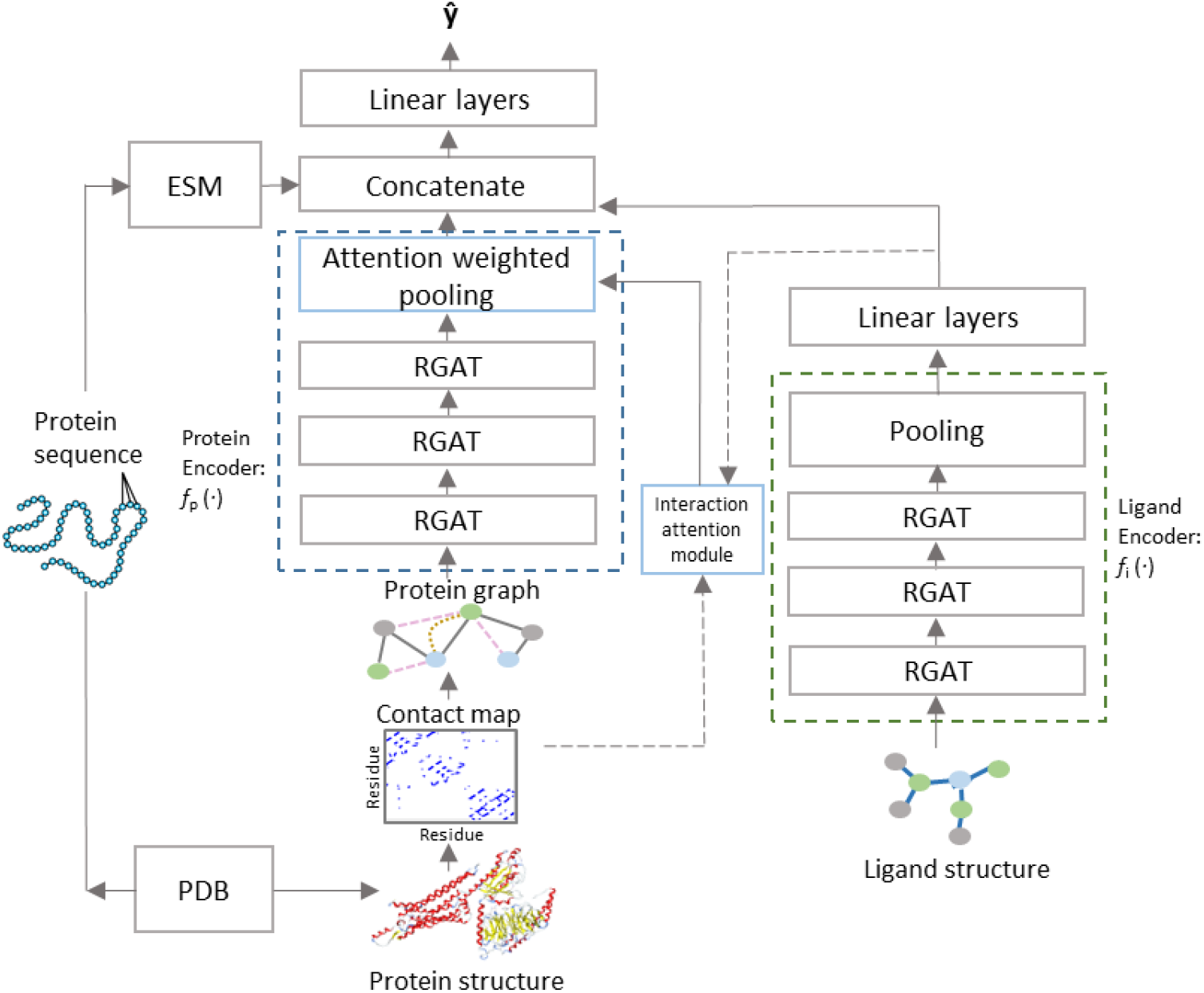
PSG-BAR Architecture

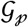 and 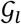 are processed by independent and architecturally identical protein and ligand encoders (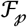 and 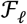 respectively) that stack several layers of GAT with skip connection. For ligand encoder 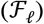, the vector representation learned by GAT layers are aggregated over all nodes of the graphs using a readout function *r*(·) that combines global max and average pooling over all nodes of the graph to generate graph level representations 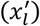. The readout function for the protein encoder performs global max pooling over the node embeddings weighted by the attention map learned by the interaction attention module. The encoded latent representations for protein are further enriched with continuous dense embedding for protein backbone, provided by a pretrained language model trained on amino acid sequences^44^. 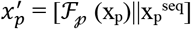 and 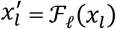. The protein and ligand representations interact with each other in a Siamese-like fusion approach (Figure 2). The decoder comprises a multilayered perceptron with LeakyReLU activation and forms the predictor function.

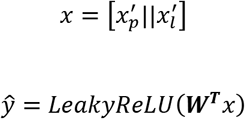

**Figure 2.**
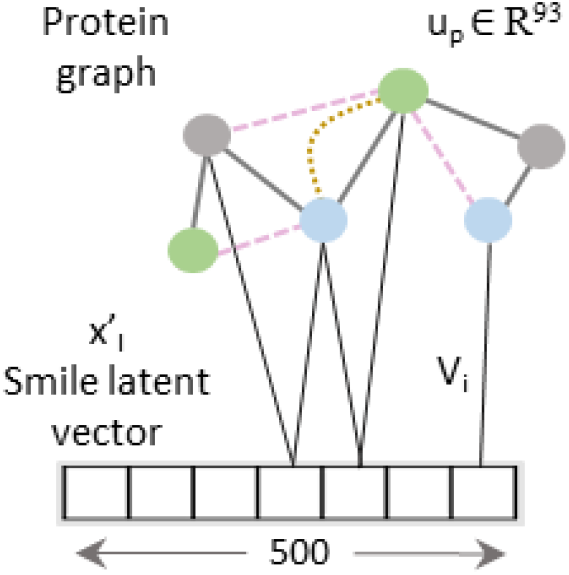
Virtual edge set to calculate cross-attention between protein graph nodes and learned representation of the drug.

### 2.3 Interaction Attention Module

To further enhance the predictive capability of our model and to learn from the relationship between protein and ligand interaction, we propose an interaction attention module based on cross attention mechanism (Figure 2). Cross attention learns to selectively attend to the nodes of the protein structure graph and hence identify principal nodes for a given protein-ligand interaction pair. We create virtual edge set 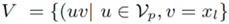 to connect all nodes of the protein graphs to the drug representation out of the ligand encoder. *V_i_* is calculated as

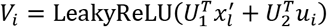

where, *U*_1_, *U*_2_ are trainable model parameters and 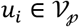

The attention map is then defined as 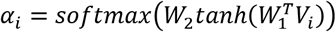 analogous to Bahdanau et al. (2014)^45^.

### 2.4 Training and hyperparameter tuning

In all our experiments we employed a base model with 3 stacked GAT layers as encoder (Figure 1). LeakyReLU activation was used throughout the model, except in the interaction attention module where *tanh* was used. All GAT layers had a dropout rate of 10% and batch normalization to avoid overfitting. Further, early stopping conditioned on validation loss was also adopted. Adam optimizer with starting learning rate of 0.07 and a decaying learning rate scheduler was employed for all our experiments. Minibatch size of 256 was found to be most suitable on our Nvidia Tesla V100 GPUs. Mean squared error and cross entropy loss were used as objectives for regression and classification respectively. All hyperparameters were empirically chosen to maximize Pearson correlation metric. Our methods were implemented using Pytorch-geometric and Pytorch.

## 3 Results

### Evaluation Metrics

Our models were evaluated for regression using Pearson coefficient and mean squared error (MSE). Additionally, for activity classification on SARS-CoV datasets, we report Cross Entropy Loss and area under the curve of Receiver Operating Characteristic curve.

#### 3.1 Binding Affinity prediction results

To test the robustness and generalizability of our method, we test the performance under following settings for each benchmark dataset (Table 2). These settings are based on the different stratification criteria of proteins and ligands for the train-test split.

**Table 2.**
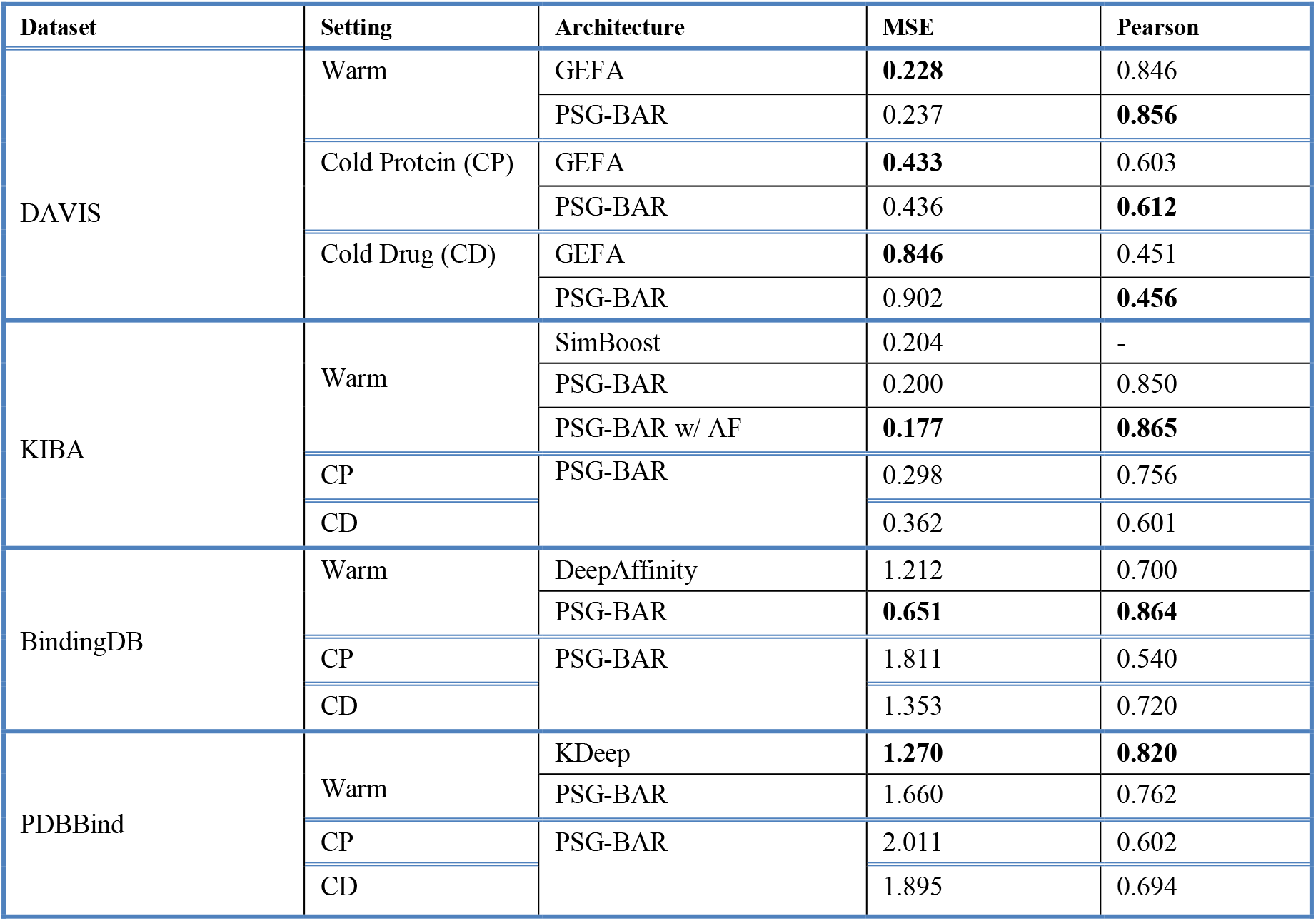
Comparison of PSG-BAR to other state-of-the-art methods on popular benchmark datasets. For each of the 4 datasets, we compare PSG-BAR to best reported performance on that dataset that we find in literature survey. For brevity, RSME data was consolidated into MSE when not directly available from the authors.

##### Warm setting

No splitting restriction imposed on proteins and ligands. Any protein (or ligand) may be repeated in training and test split; however, interactions are not duplicated across the 2 splits.

##### Cold Protein setting

Each unique protein in the dataset has restricted membership to either training or test set. No restrictions on ligands.

##### Cold Ligand setting

Each unique drug in the dataset has restricted membership to either training or test set. No restrictions on proteins.

#### 3.2 SARS inhibitors prediction

To perform a binary classification on SARS-CoV Mpro inhibitor and to overcome the problem of class imbalance between actives and inactives, we oversampled the actives and randomly subsampled inactives to construct a class-balanced dataset of 26,400 interactions. With a train-test split of 80-20% and 5-fold CV, our method yielded 0.72 ROC-AUC (Figure 3A). The sparsity of actives within an expansive and diverse chemical space warrants identification of even slightly probable hit compounds for further evaluation. As such, we optimize for recall by using weighted binary cross entropy loss.

**Figure 3.**
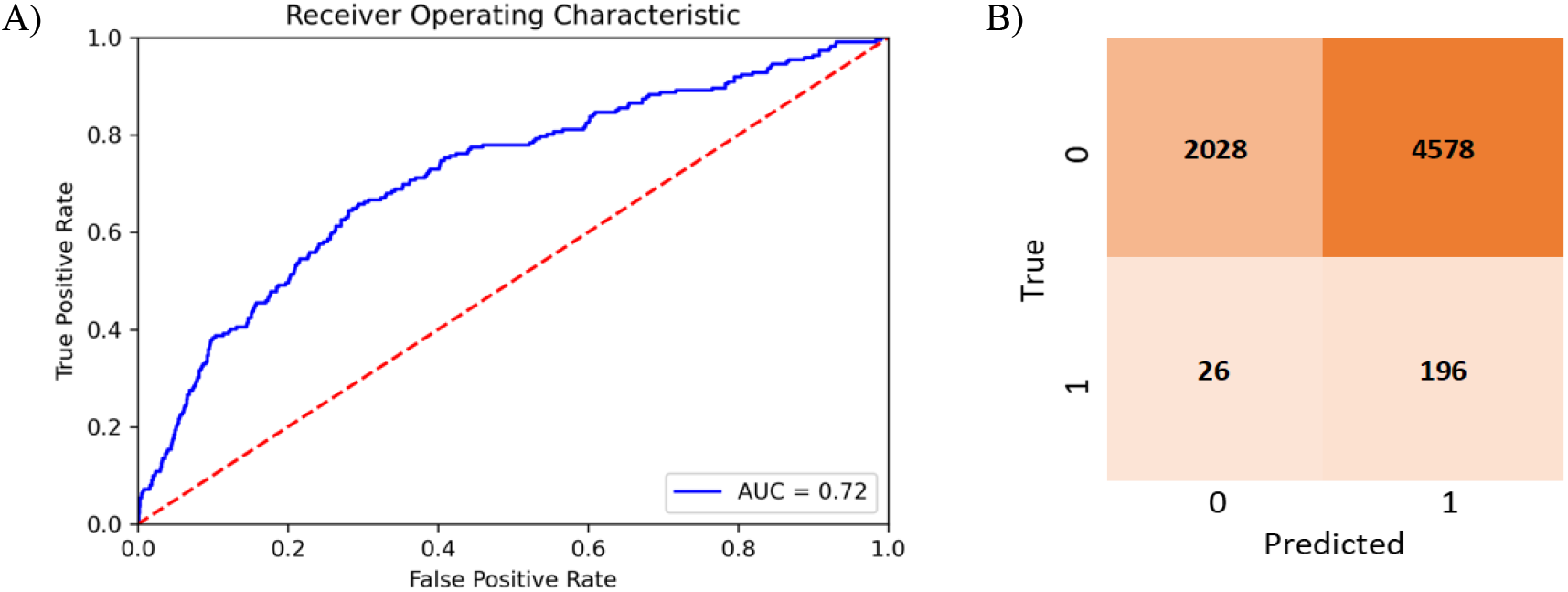
PSG-BAR predictions for SARS-CoV inhibitors A) ROC plot B) Confusion matrix 0: inactives; 1: actives.

#### 3.3 SARS CoV-2 MPro experimental validation

To validate the utility of PSGBAR method, we additionally perform testing on the novel coronavirus, SARS-CoV-2, the causative agent of the ongoing COVID-19 global pandemic. We rank PSG-BAR scores against docking screen results for Mpro target. This docking screen^46^ was performed on the Mpro crystal structure (6W63^47^) using DeepDocking^7^ tool coupled with commercial software Glide^48^. Importantly, after the docking ranking, the final set of 1200 molecules were selected by expert chemists. These selected compounds were then experimentally tested and 116 actives were confirmed. We observe that our model highly ranks the compounds that the human expert decided are worth purchasing. We show the normalized histograms of predicted scores for docking screen compounds (blue) and decoys (orange). The details of our findings are as following:

1. The predicted score distribution for hits from docking is right skewed, meaning that the model highly ranks most compounds in agreement with the docking program’s ranking.
2. The flat, near-zero left tail indicates that the tested set has almost no compounds which are predicted to be extremely poor-binders by our method. This aligns with our *a priori* knowledge that these tested compounds are expert selected following docking screen.
3. A set of decoys of a random sample of 20,000 poorly ranked compounds by docking yield low score on our model too. Figure 4 illustrates that on average, hit compounds exhibit higher PSG-BAR scores than non-hits. This corroborates the utility of our method as a standalone method to screen large virtual libraries. However, the most lucrative positioning of our method would be as a filter succeeding docking and preceding hit compounds purchase.

**Figure 4.**
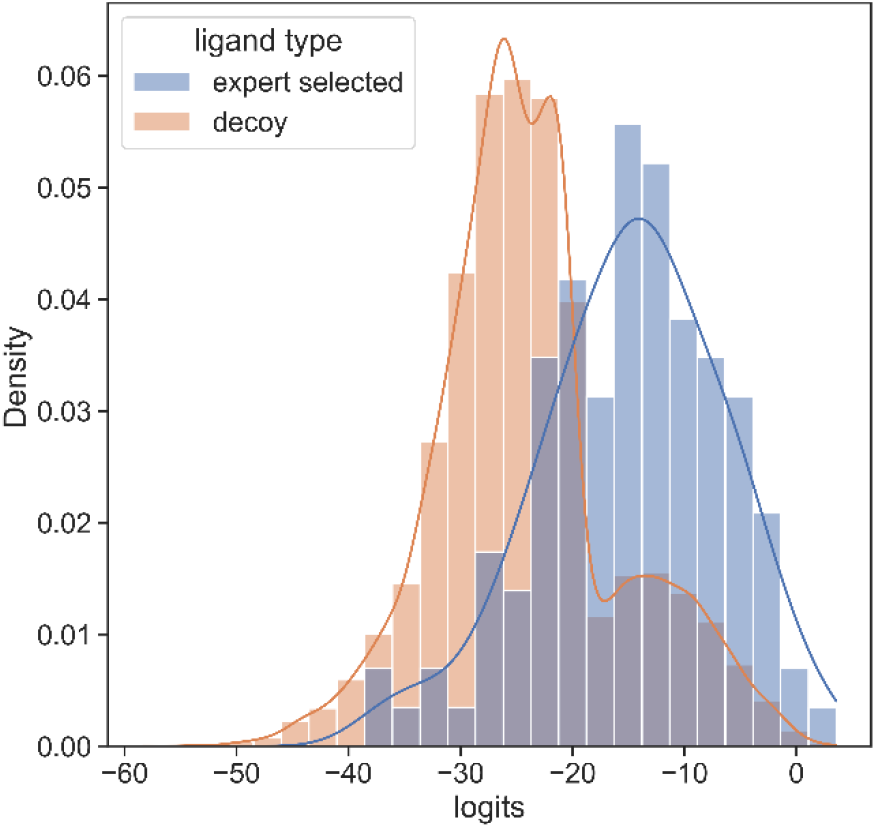
Histogram of PSG-BAR scores for docked compounds suggests that our method assigns higher scores hits from docking and low scores to poor binders. The distribution to the right shows the PSG-BAR scores for top ranking hits of docking screen for MPro target. The left distribution is of PSG-BAR scores for randomly chosen 20,000 compounds from docking screen that were not considered as hits. These compounds on an average had lower PSG-BAR score (−30) compared to scores of hit compounds (−15). PSG-BAR scores are unnormalized logits of our trained deep neural network.

#### 3.4 Attention Centrality

Thus far, we have evaluated the performance of our model as a whole on several datasets and use cases. We are further interested in investigating the interpretability of method’s predictions and understanding the causality for its superior performance. The relevance of the attention scores was evaluated on a randomly selected sample of 50 proteins from six protein families namely, protease, kinase, polymerase, transferase, phosphodiesterase and hydrolase from PDBBind dataset. We focus on the amino acid residues with the highest attention scores, which should be relevant for predicting binding. Figure 5 illustrates the distributions of average Solvent Accessible Area (SAA) for all protein residues and the top-10 residues with the highest learned interaction attention scores. Notably, residues with the highest attention scores displayed higher mean SAA for all protein families. We also observed that, on an average, surface residues have 22% higher attention scores than core residues. This aligns with our domain knowledge on protein ligand binding that suggests the pocket residues located on the protein surface or are usually somehow accessible to the solvent.

**Figure 5.**
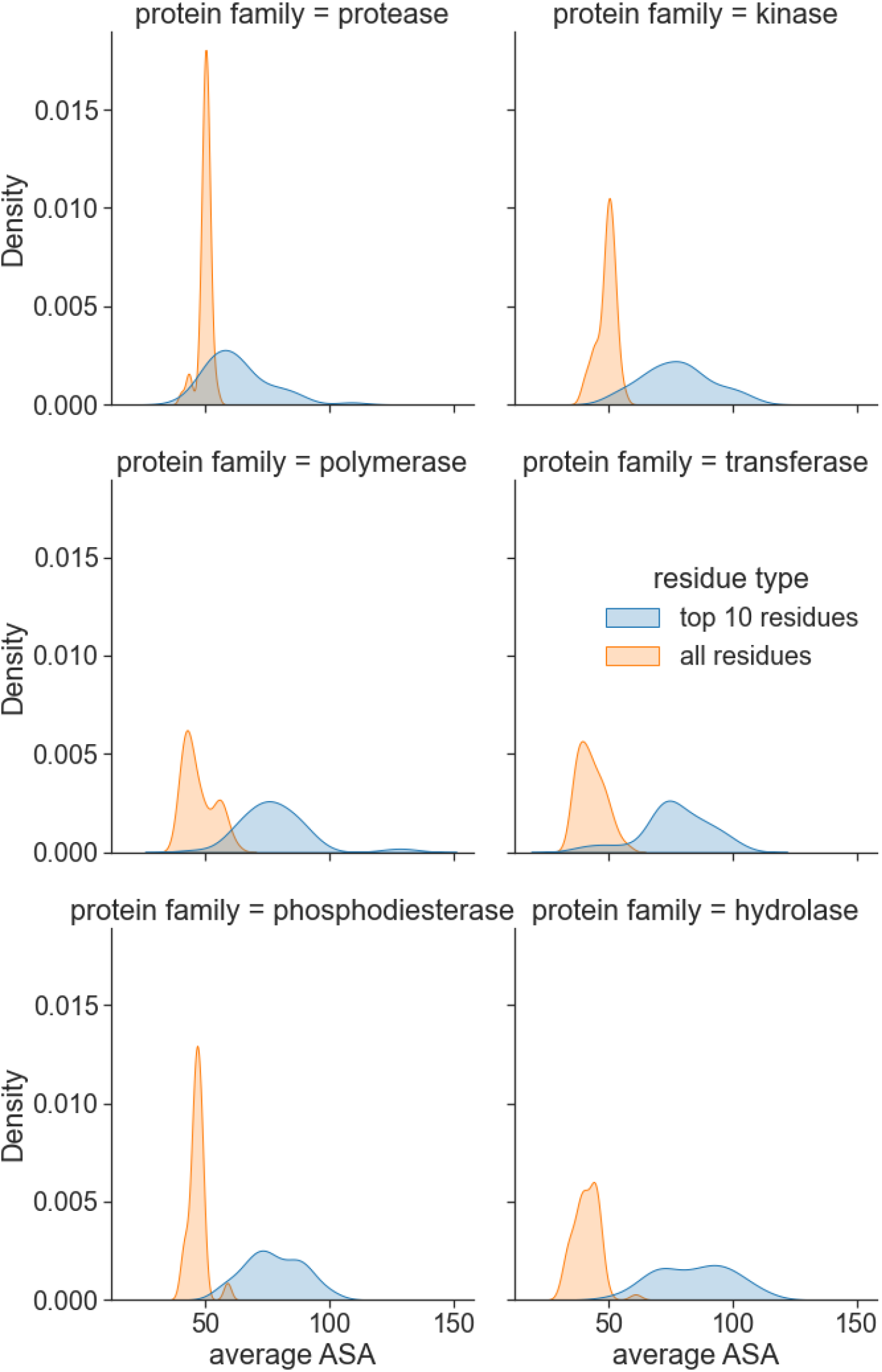
Interaction Attention scores predicts residues on the protein surface. For the 6 most frequent protein families in PDBBind dataset, highest scored residues by PSG-BAR have mean SAA higher than that of the remaining residues.

#### 3.5 Drug Promiscuity

PSG-BAR operates as a protein target agnostic method for ligand binding prediction. As such, it could be utilized to study the off-target effects of drug compounds on a multitude of human proteins. To this end, we evaluate the promiscuity of the hit compounds discovered in section 3.3. To train a model that predicts activity across major off-target proteins, we used a dataset presented by Ietswaart et al (2020)^49^ that contains *in vitro* activity of 1866 marketed drugs across 35 protein targets (38,091 protein-ligand interactions). These targets are adverse-drug reaction (ADR) related and include well-known proteins such as hERG (induces cardiotoxicity), nuclear receptors (carcinogenic effects) and COX enzymes (intestinal bleeding). Thus, our model predicts whether a molecule is likely to bind any of these off-target proteins. The modified PSG-BAR method used to perform classification for activity on ADR dataset yields ROC-AUC value of 0.88 (Figure 6A).

**Figure 6.**
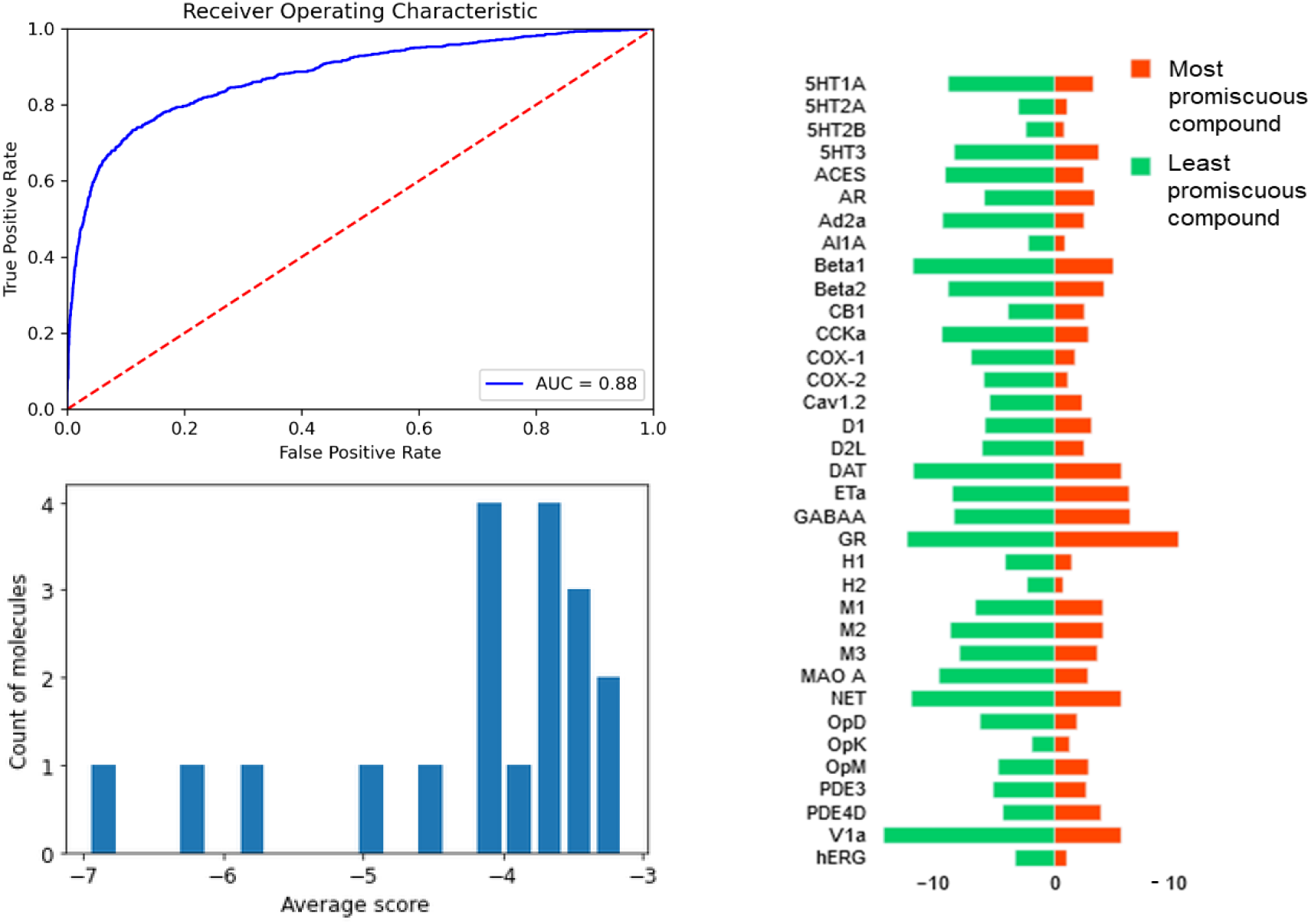
Evaluation of off-target effects of MPro hits on human proteins using PSG-BAR. (A) ROC-AUC on ADR dataset (B) Histogram of average scores across 35 ADR-related proteins for 19 predicted Mpro binders. The average score is a proxy for compound promiscuity. (C) Predicted likelihood of binding to ADR proteins for the most promiscuous molecule (red) and least promiscuous molecule (green). Lower values (closer to the center) indicate a high likelihood of binding.

To evaluate effects on ADR proteins, we chose 19 compounds ranked within the top 25^th^ percentile by both PSG-BAR and docking screen. These compounds are predicted to be the most likely binders to Mpro by our pipeline. We averaged scores across all proteins to assess general promiscuity of a molecule, i.e. the likelihood of binding to many proteins. The average score ranged from −3 to −7, where higher score means more promiscuous compound (Figure 6B). We examined two molecules from both extremes. The most promiscuous molecule was predicted to most likely bind alpha-1A adrenergic receptor (Al1A), histamine H2 receptor (H2), COX-2, hERG and 5-hydroxytryptamine receptor 2A (5T2A) and 2B (5HT2B) (Figure 6C). Thus, if this molecule in the process of drug development turned out to be problematic (e.g. exert toxicity in mice), our model can hint to the possible underlying off-targets. Such information might be extremely useful for structure-based optimization of the lead compound.

#### 3.6 Ablation Studies

We critically analyze the contribution of key components of our model towards predictive performance. These studies were conducted on the PDBBind dataset under warm setting, train/validation split of 0.8/0.2 ratio and with a fixed random seed to impose fair uniformness across experiments. We first study the effect of the skip connections in GATs. The protein and ligand encoders are equipped with varying number of GAT layers while iteratively ablating the skip connections for each constituent layer.

We observe that stacking graph layers has diminishing returns (Figure 7A). This is consistent with the observations made by previous works as most state-of-the-art GNNs are shallower than 4 layers. Our experiments show similar peak performance for 3-layered network. In each of the stacking mode, GAT network equipped with the skip connection performed better than the corresponding non-residual GAT version. Further stacking layers reduce the overall efficacy of our model however it continues to benefit from the addition of the skip connection. Furthermore, we noticed that over the early epochs of model training, non-residual GAT yields slightly better performance than skip-connection variant and converges faster (Figure 7B). This may be attributed to (i) comparatively simpler architecture of the former and (ii) averaging of node representations to nearly same vector to cause oversmoothing occurs over the due course of passes over the data.

**Figure 7.**
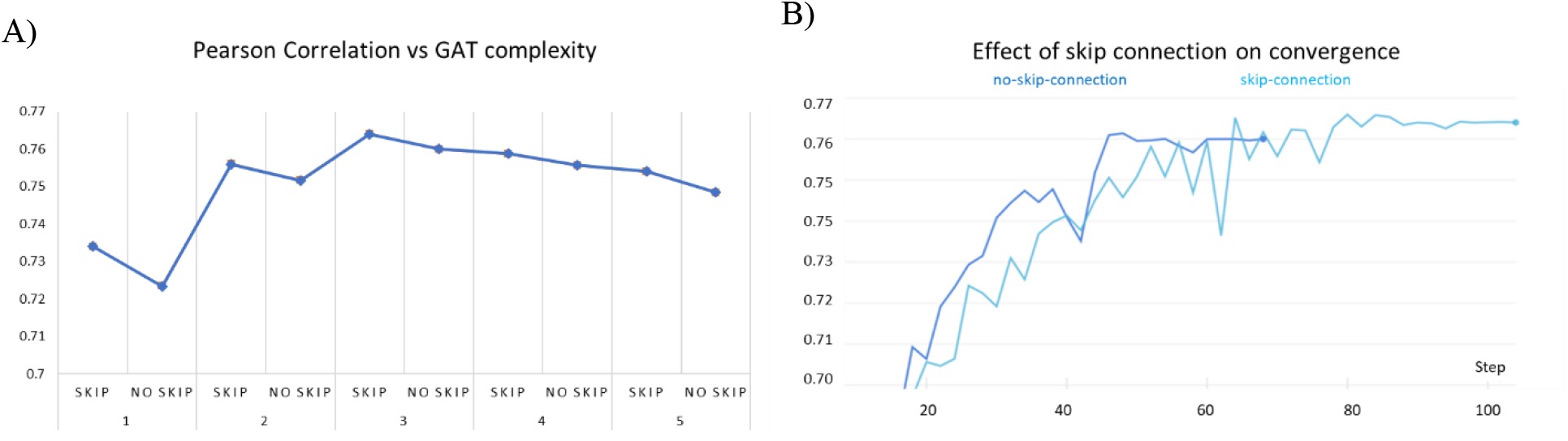
Effect of skip-connection on model performance on PDBBind dataset. (a) skip-connection on stacking GAT layers compared to GAT models of same complexity without skip connection. (b) skip connection vs no skip connection on 3 layered GAT: the early success of no-skip variant is superseded by skip-connection variant over increasing epochs.

We further investigated the effect of the pre-trained amino acid embeddings on the overall performance. In the model without amino acid embeddings, we observed a 3.47% decrease in the Pearson correlation and 4.93% increase in the MSE on the PDBBind dataset. This is attributed to the fact that protein homology is driven by sequence similarity and a well-trained embedding on large amounts of amino acid sequences effectively captures such protein similarity.

##### 3.6.1 Effect of augmenting KIBA dataset with AlphaFold structures

As described in dataset section, utilization of any protein-ligand interaction in the training (or validation) process was contingent upon the availability of the crystallographic structure of the corresponding protein target. For the KIBA dataset we supplemented the proteins with missing PDB with their corresponding structures from AlphaFold. Consequently, the additional interaction samples led to a 43.4% increase in available proteins in the usable dataset and a 19.67% increase in total interactions. The model performance also rose by 1.72% in terms of Pearson correlation and 11.1% for MSE (table 2).

##### 3.6.2 Effect of secondary structure features of proteins

We hypothesized that additional protein structure descriptors derived from DSSP module will help to ameliorate the definition of binding pockets. Indeed, Stank et al. highlighted examples where secondary and tertiary features aided the classification of various protein pockets^50^. Furthermore, topological, solvation and hydrophobicity descriptors may help determine the druggability of protein sites. For instance, it is known that most druggable sites are highly hydrophobic and relatively deep^51^. As expected, the aforementioned features boosted the performance of PSG-Bar with Pearson correlation increase in the range of 1.3-1.76% and MSE decrease in the range of 4.00% to 4.81% (Table 3). This could indicate the benefit of implicit learning of the connection between properties of amino acid residues in the pocket and the binding free energy of ligands.

#### 3.7 Error analysis of prediction of effective binders

It should be noted that drug discovery practitioners tend to care more for prediction on good binders rather than for the overall performance of computational approaches. To this extend, we stratify the affinity score range into 4 intervals as <=10, 10-12, 12-14 and >14 for the KIBA dataset (Figure 8). A significantly large majority (78.4%) of interactions lie in the score 10-12 interval. These are weak binders, and the model performance is at its peak for these interactions. The MSE for this stratum is 0.093 compared to the population mean MSE of 0.2. Further, the moderate binders (score 12-14) and strong binders (score > 14) span much smaller proportion of total interactions at 18.4% and 2.7% respectively. For the poor binders, the predictive performance is considerably inferior to any other strata.

**Figure 8:**
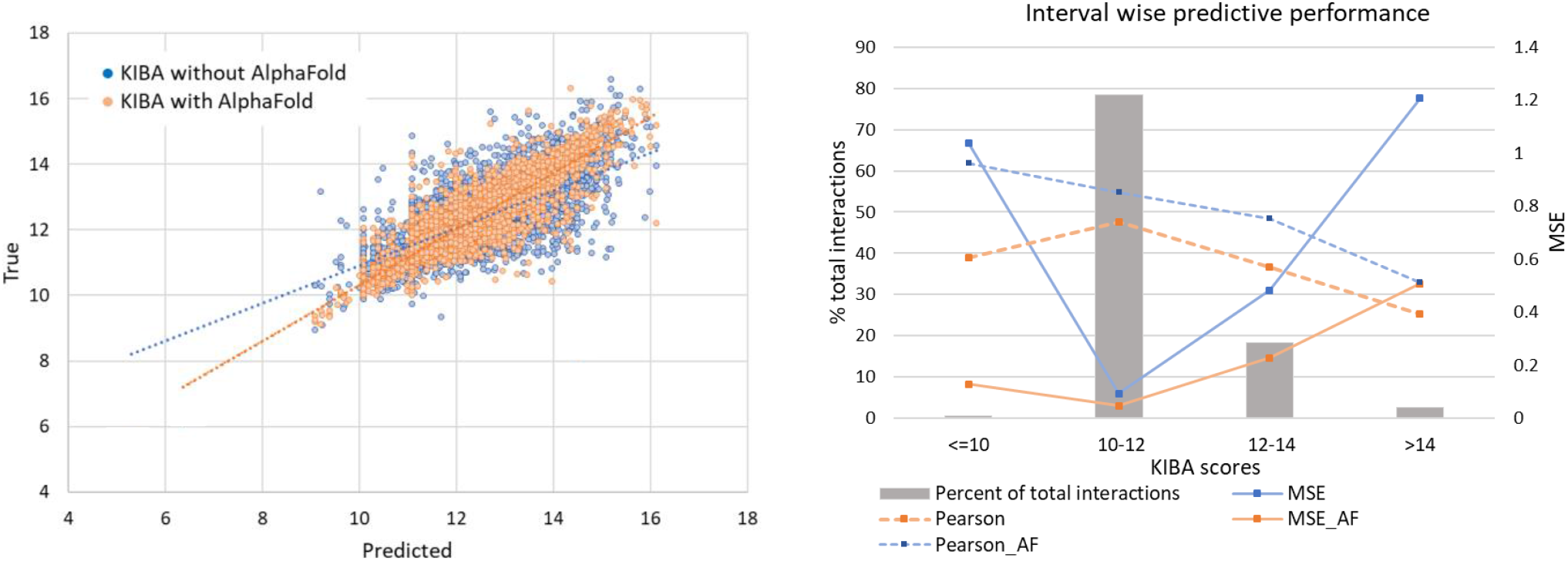
A) Linear correlation of experimental vs predicted pKd values from the KIBA dataset. B) Interval wise predictive performance on KIBA dataset. The most common interval of 10-12 has highest performance while the extremes on both sides have very poor performance.

It should be mentioned however that these interactions are of relatively low significance to design and discovery of drugs. Strong binders (score > 14) are unarguably the interval of highest interest and difficulty to predict. This, coupled with their sparsity in the evaluated subset of the data, leads to a mediocre result for strong binders (MSE 1.2 compared to MSE 0.2 overall). However, further elaborating the KIBA dataset with AlphaFold structures, show a marked improvement of 57.9% in MSE to bring down MSE to 0.50. This is due to the presence of more structurally similar proteins that enables the model to learn complex binding features, central to strong protein-ligand binding.

## Discussion

In this study we report the development of PSG-BAR, a ligand binding affinity prediction method leveraging protein 3D structures, sequence embedding, and 2D graph representation of a drug molecule. Many reported studies concerning protein-ligand interaction fail to consider the complex folded 3D structure of the proteins and employ just the primary protein sequence. Yet others that consider protein 3D structure, fall short to consider the relevance of active sites of the protein and view all the protein residues as equally important. Some other studies utilizing binding site information rely on computationally expensive methods to determine binding sites apriori and hence restraining their applicability to diverse datasets. PSG-BAR alleviates these limitations by using entire protein structural graph and learning attention scores to selectively weight useful regions of the protein based on its interaction with the drug molecule. As a result, our method outperforms state-of-the-art affinity prediction methods on several benchmarking datasets.

As such, the integration of protein structures helps to achieve better predictive results. This is mainly because 3D structures contain relevant information on actual configuration of the binding pockets, which have immediate implications for the ligand binding. These methods are bottlenecked by the availability of experimentally derived protein structures; however, with the advancement of NMR Xray crystallography and cryo-EM techniques, more high resolution PDBs are being deposited than ever before. Furthermore, as a result of Alphafold, even more predicted protein structures became available. These developments enable effective advancement of deep learning based approaches; in this work we validate this hypothesis by predicting experimentally determined measures of binding affinity on several protein targets across standard benchmark datasets. Particularly for the KIBA dataset, we show that the augmentation with Alphafold structures improves MSE by 11.1%.

It should also be emphasized that augmentation of 3D protein structure information with 2D sequence descriptors can further improve model performance. Since protein sequences capture some level of structural similarity, and are relatively easily accessible than the crystallographic structures, they become excellent candidates to learn large-scale protein embedding using unsupervised pretraining. As such, enriching our model with a sequence embedding trained on 250M protein sequences yields better generalizability for *warm-protein setting* across all tested datasets.

The training on the abundance of diverse proteins makes the model highly generalizable across different protein families. Thus, the model could be applied for predictions across a set of selected protein such as to ADR-relevant targets. Such a model could be used to evaluate if the selected drug candidates are promiscuous and have toxic off-target effects. PSG-BAR is target agnostic and provides a reliable means for such off-target analysis. We evaluated the selected Mpro lead compounds using our model built for prediction across ADR-related proteins. We found that the molecules range in their predicted promiscuity. Thus, these predictions might help to guide future lead optimization of the drug candidates.

We acknowledge other works with better reported MSE especially on the PDBBind dataset (KDeep). This gain is largely attributed to the utilization of protein-ligand complexes in their model indicating that the binding pockets (active sites) of the proteins are most critical towards this downstream prediction. However, obtaining such protein-ligand complex is expensive and hence limits the applicability of many of these methods to smaller datasets with selective protein targets. Similarly, conventional docking approaches depend on computationally expensive preparation of binding sites. Our attention-based method learns surface residues without direct supervision while simultaneously predicting binding affinity. This identification of key binding residues diminishes the need for expensive binding site preparation and makes our model accessible to minimally preprocessed protein structures.

In conclusion, our work represents a step in the direction of alleviating the problem of *a priori* knowledge of binding site which is an expensive pre-requisite to all protein-ligand interaction studies. The road to explainable AI, such as in attention based visual question answering, is expected to reform deep learning for DTI. We expect more protein-ligand complexes to be experimentally resolved, so next generation of deep models will be able to learn such attention scores more accurately. In fact, with soft-supervision these attention scores may even lead to reliable identification of binding sites. In this study, we work under the rigidity assumption of protein and ligand (by using only 2D molecular structures). Further research should investigate 3D ligand conformers in conjunction with flexible protein 3D structures.

## Acknowledgement

This work was funded by Canadian Institutes of Health Research (CIHR) (AWD-015014), Canadian 2019 Novel Coronavirus (2019-nCoV) Rapid Research grants (OV3-170631 and VR3-172639), and generous donations for COVID-19 research from TELUS, Teck Resources, 625 Powell Street Foundation, Tai Hung Fai Charitable Foundation, Vancouver General Hospital Foundation. The authors thank the Dell Technologies HPC and AI Innovation Lab for their support and partnership in providing the HPC platform (PowerEdge servers) to accelerate the AI algorithms, and the UBC Advanced Research Computing team for providing access and technical support for the Sockeye supercomputing cluster.

---

**Table S1:** Protein surface features improves PSG-BAR predictions across all 4 benchmarked datasets.

|  | With Surface Features |  | Without Surface Features |  |
| --- | --- | --- | --- | --- |
| Dataset | MSE | Pearson | MSE | Pearson |
| BindingDB | 0.651 | 0.864 | 0.678 | 0.851 |
| PDBBind | 1.660 | 0.762 | 1.744 | 0.749 |
| KIBA | 0.200 | 0.850 | 0.209 | 0.837 |
| DAVIS | 0.237 | 0.856 | 0.249 | 0.845 |

